# Industrialization of three-dimensional hiPSC-cardiac microtissues for high-throughput cardiac safety and drug discovery screening

**DOI:** 10.1101/2024.11.29.626032

**Authors:** Tessa de Korte, Benjamin B. Johnson, Georgios Kosmidis, Benoit Samson-Couterie, Mervyn P. H. Mol, Ruben W. J. van Helden, Louise François, Viviana Meraviglia, Loukia Yiangou, Tom Kuipers, Hailiang Mei, Milena Bellin, Stefan R. Braam, Shushant Jain, Christine L. Mummery, Richard P. Davis

## Abstract

Current cardiac cell models for drug screening often face a trade-off between cellular maturity and achieving high throughput. While three-dimensional human induced pluripotent stem cell-based heart models typically exhibit more adult-like features, their application is hindered by the need for large cell numbers or complex equipment. Here, we developed cost-effective methods to scale up production of three-dimensional cardiac microtissues (cMTs) containing three cardiac cell types, and assess calcium transients and action potential metrics for high-throughput screening (HTS). Automating the procedure revealed reproducible drug responsiveness and predictive accuracy in a reference compound screen. Furthermore, an arrhythmic phenotype was reliably triggered in cMTs containing cardiomyocytes with a RYR2 mutation. A screen of FDA-approved drugs identified 17 drugs that rescued the arrhythmic phenotype. Our findings underscore the scalability of cMTs and their utility in disease modelling and HTS. The advanced “technology-readiness-level” of cMTs supports their regulatory uptake and acceptance within the pharmaceutical industry.

## Introduction

The use of human induced pluripotent stem cell-derived cardiomyocytes (hiPSC-CMs) is becoming pivotal in drug discovery and development as evidenced by their regulatory recognition, which includes FDA and EMA acceptance for nonclinical cardiac safety risk assessments and their inclusion in the FDA Modernization Act for testing drug effectiveness^1,2^. Nonetheless, the ability of hiPSC-CMs to reveal disease phenotypes and assess how compounds affect the heart, could be enhanced by furthering their maturity to resemble adult rather than foetal states^3,4^. Recent developments in hiPSC-derived three-dimensional (3D) cardiac models have shown to improve cardiomyocyte maturity^5–10^. Nevertheless, these models often require specialized knowledge, substantial cell numbers or specific equipment^11^, rendering them costly and making them challenging for routine implementation in high-throughput screenings (HTS). These limitations are compounded by the need to ensure scalability, compatibility with existing assays and reproducibility; key criteria for pharmaceutical industry adoption^12^.

Furthermore, for these 3D models to be effective in drug efficacy screening, they must consistently and accurately replicate specific disease phenotypes^13,14^. For instance, catecholaminergic polymorphic ventricular tachycardia type 1 (CPVT1) is an inherited arrhythmogenic disorder associated with over 170 mutations distributed throughout the ryanodine receptor 2 gene (*RYR2*)^15^. This diversity in mutations, combined with incomplete penetrance and variable expressivity, results in a wide range of phenotypes and disease severities, complicating diagnosis and treatment^15,16^. Previous studies using hiPSC-CMs to investigate CPVT1 disease mechanisms have shown their potential to reveal disease-associated phenotypes, such as susceptibility to stress-induced arrhythmias, and to recapitulate drug responses^17^. However, these studies have faced challenges with throughput and the reproducibility of revealing the arrhythmic phenotype^18–23^, thus limiting their suitability for HTS. Thus, there is a pressing need for hiPSC-cardiac (disease) models that are scalable, cost-effective, and consistently display a reproducible phenotype suitable for HTS of new or repurposed drugs.

Here, we established pipelines for scalable, affordable, and scaffold-free generation of cardiac microtissues (cMTs)^7^ from cryopreserved batches of hiPSC-CMs, - cardiac endothelial cells (cECs), and -cardiac fibroblasts (cFbs). These cMTs supported fluorescence-based calcium (Ca^2+^) and voltage assays for cardiac safety and efficacy assessment, demonstrated reproducibility across multiple cell batches, and accurately reflected known responses to cardioactive drugs. By incorporating hiPSC-CMs containing a RYR2 mutation in the cMTs and subsequently treating these with the membrane-permeable dibutyryl cAMP (dbcAMP), we established a robust model of the CPVT1 arrhythmic phenotype. This phenotype was consistently rescued using flecainide, a cardiac sodium- and RYR2-channel blocker that has emerged as an effective secondary anti-arrhythmic agent in patients suffering from CPVT1^24,25^. Robotics were employed to automate the formation, maintenance, and dye labelling of cMTs. Using this approach, blind screens of reference compounds with known inotropic effects validated the accuracy and reproducibility of the cMTs and Ca^2+^ assays. Subsequently, a larger screen of 92 FDA-approved drugs identified 17 compounds that rescued the arrhythmic phenotype associated with CPVT1.

Our study demonstrates that 3D cMTs are scalable, automatable, and offer a predictive approach for HTS. High reproducibility, proper validation and standardization as shown here are essential steps towards pharmaceutical industry adoption and regulatory authority acceptance of these models.

## Results

### Standardisation of cardiac microtissue generation and establishment of optical calcium and voltage readouts

To refine cMT generation, we modified the initial step for differentiating hiPSCs into cardiac mesoderm. Instead of using the cytokines BMP4 and Activin A^26^, we adopted a protocol that exclusively utilises the Wnt-signalling pathway activator CHIR99021, in line with established hiPSC-CM differentiation procedures^26–28^. This “small molecule protocol” (SP) adaptation (**Supplementary Fig. 1a**) generated mesoderm that could also subsequently be differentiated to cardiac endothelial cells (cECs) and epicardial cells (EPCs), the latter being a precursor of cardiac fibroblasts (cFbs)^7^. The EPCs exhibited typical epithelial cobblestone-like morphology (**Supplementary Fig. 1b**) and expressed the transcription factors WT1 and GATA4, and the gap junction protein, connexin 43 (CX43), while the resulting cFbs lost WT1 expression (**Supplementary Fig. 1c**). The SP resulted in >80% of the differentiated cells expressing markers associated with CMs (cardiac troponin T (cTnT))-, cECs- (platelet and endothelial cell adhesion molecule 1 (CD31)) or cFbs (vimentin) across multiple differentiations, indicating a high degree of standardization and cell type purity (**Supplementary Fig. 1d**). RNA sequencing was performed to compare these cells with those derived from the cytokine protocol (CP) and respective primary human tissue equivalents^7^, with principal component analysis demonstrating close clustering of hiPSC-derived cell types regardless of the differentiation protocol (**Supplementary Fig. 1e**). Interestingly, SP-derived cECs appeared to cluster more closely with primary ECs from later-stage foetal heart samples (week 15 and week 21), and showed less variance compared to CP-derived cECs. Based on these observations, we decided to use SP-derived cardiac cells for cMT formation.

Cryopreserved stocks of the cardiac cell types were used to form cMTs, which self-assembled into contracting spheroids within 48 h. After 2 weeks in culture, these cMTs had a similar morphology (diameter: 425.3 ± 74.4 µm) and cellular architecture (**Fig 1a**), to that previously reported^7^. To assess cMT functionality, we developed fluorescence-based assays for evaluating Ca^2+^ handling and voltage kinetics in a high-throughput format capable of measuring up to 384 cMTs simultaneously. The cMTs displayed robust signals with low variability in beat rate, amplitude and peak width duration at 50% decay (PWD50), and an intra-well peak-to-peak variability of 5% in Ca^2+^ and 14% in voltage measurements (**Fig. 1b**). Although differences in the beat rate were observed depending on whether the cMTs were labelled with a Ca^2+^ or voltage-sensitive dye, this was likely because of the ion-chelating properties of the dyes on contractile behaviour of cardiomyocytes^29^.

**Figure 1.**
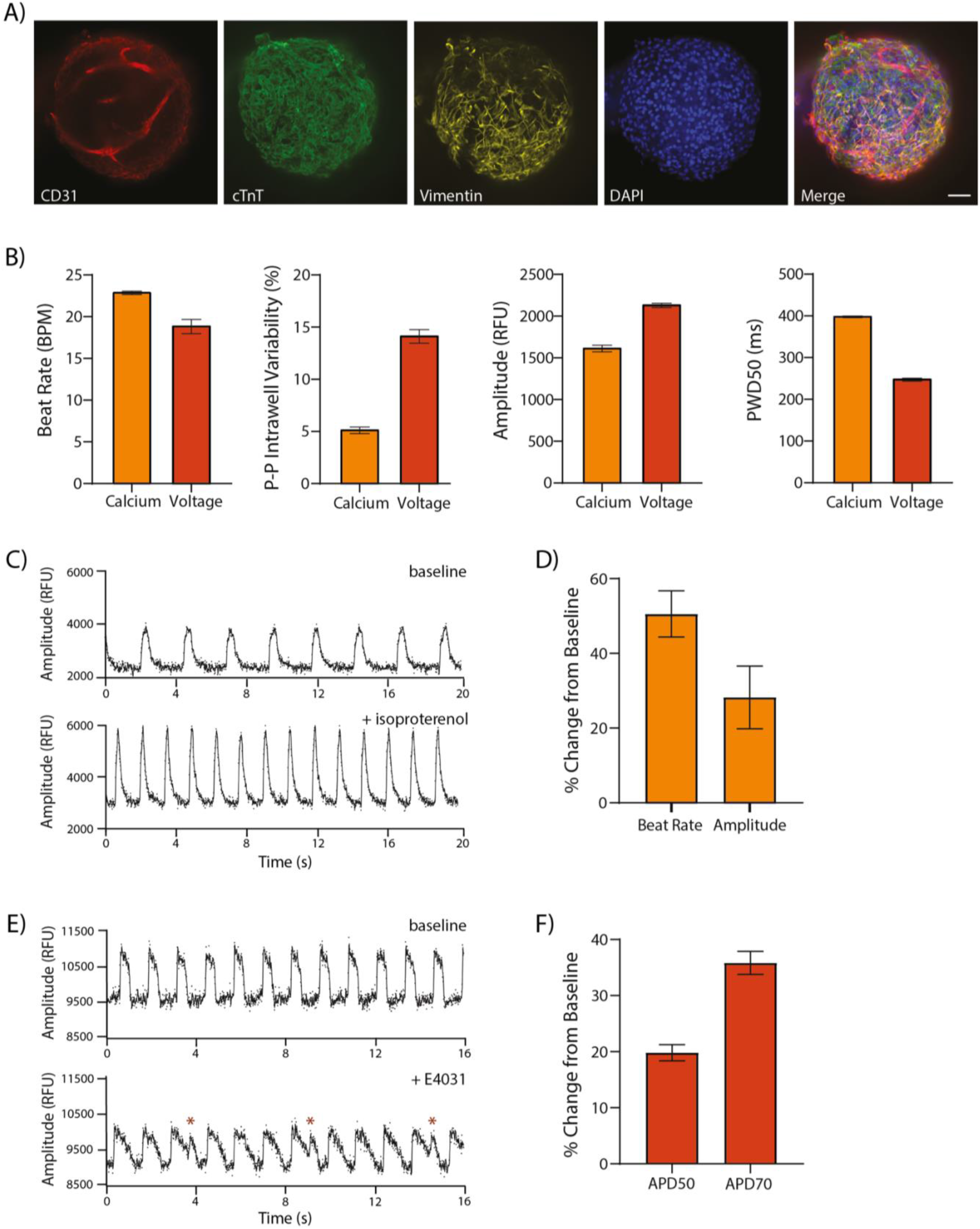
Evaluation of Ca^2+^ Handling and Voltage Kinetics in cMTs. **A)** Representative immunofluorescence images of a cMT with hiPSC-cECs marked by CD31 (red), hiPSC-CMs by cTnT (green), and hiPSC-cFbs by Vimentin (yellow). Nuclei are stained with DAPI (blue); scale bar: 100 µm. **B)** Beat rate, peak-to-peak (p-p) intra-well variability, peak amplitude, and peak width duration at 50% of decay (PWD_50_) of Ca^2+^ transient (orange) and voltage kinetics (action potential, red) in cMTs. Data is mean ±SEM; n=288 (Ca^2+^) and n=174 (voltage) from 3 independent experiments; **** p <0.0001 (unpaired t test with Welch’s correction). BPM: beat per minutes; RFU: relative fluorescent unit. **C and D)** Representative Ca^2+^ transients (**C**) recorded from a cMT before (baseline) and after addition of 0.1 µM isoproterenol, and the corresponding mean percentage change (**D**) in beat rate and Ca^2+^ peak amplitude, corrected for vehicle control (DMSO); error bars ±SEM; n=8 cMTs from 2 independent experiments. **E and F)** Representative action potential traces (**E**) recorded from a cMT before (baseline) and after addition of 0.03 µM E4031, and the corresponding mean percentage change (**F**) on action potential duration at 50% (APD_50_) and 70% (APD_70_) of repolarization, corrected for vehicle (DMSO); error bars ±SEM; n=7 cMTs from 2 independent experiments.

To validate the assays, functional testing with reference compounds was performed (**Fig. 1c-f**). Analysis of Ca^2+^ transients showed an increase in Ca^2+^ peak amplitude and beat rate 30 min after treating the cMTs with isoproterenol, a β-adrenergic agonist that has a positive chronotropic and inotropic effect. Likewise, treatment of cMTs with E4031, a hERG-channel blocker, resulted in an expected prolongation of the action potential (AP) duration and subsequent early afterdepolarization (EAD)-like arrhythmic events in the APs. These results confirmed the physiological responsiveness and suitability of cMTs for larger-scale drug screening applications.

### Automation of cardiac microtissue formation, maintenance and labelling

To facilitate the necessary upscaling of cMTs for HTS applications, we focused on minimizing manual handling by automating the cMT production and screening processes. An automated liquid handling workstation was used to streamline cMT formation, medium refreshments, and dye labelling for subsequent functional analysis (**Fig. 2a**). After optimizing variables such as cell density, mixing speed, and aspirating and dispensing parameters, the cMTs showed comparable morphology and functionality to those seeded manually. Automated cMTs were slightly smaller, with an average diameter of 373.9±37.0 µm at day 14 compared to 425.3±74.4 µm for manually seeded cMTs, but exhibited less variability (coefficient of variation (CV) ≤ 10% vs. CV ≤ 18%), indicating more consistent formation (**Fig. 2b**). The automated cMTs showed similar peak-to-peak variability in Ca^2+^ transients compared to manually seeded cMTs (**Fig. 2c**), and both responded similarly to the L-type Ca^2+^ channel blocker nifedipine (**Fig. 2d**).

**Figure 2.**
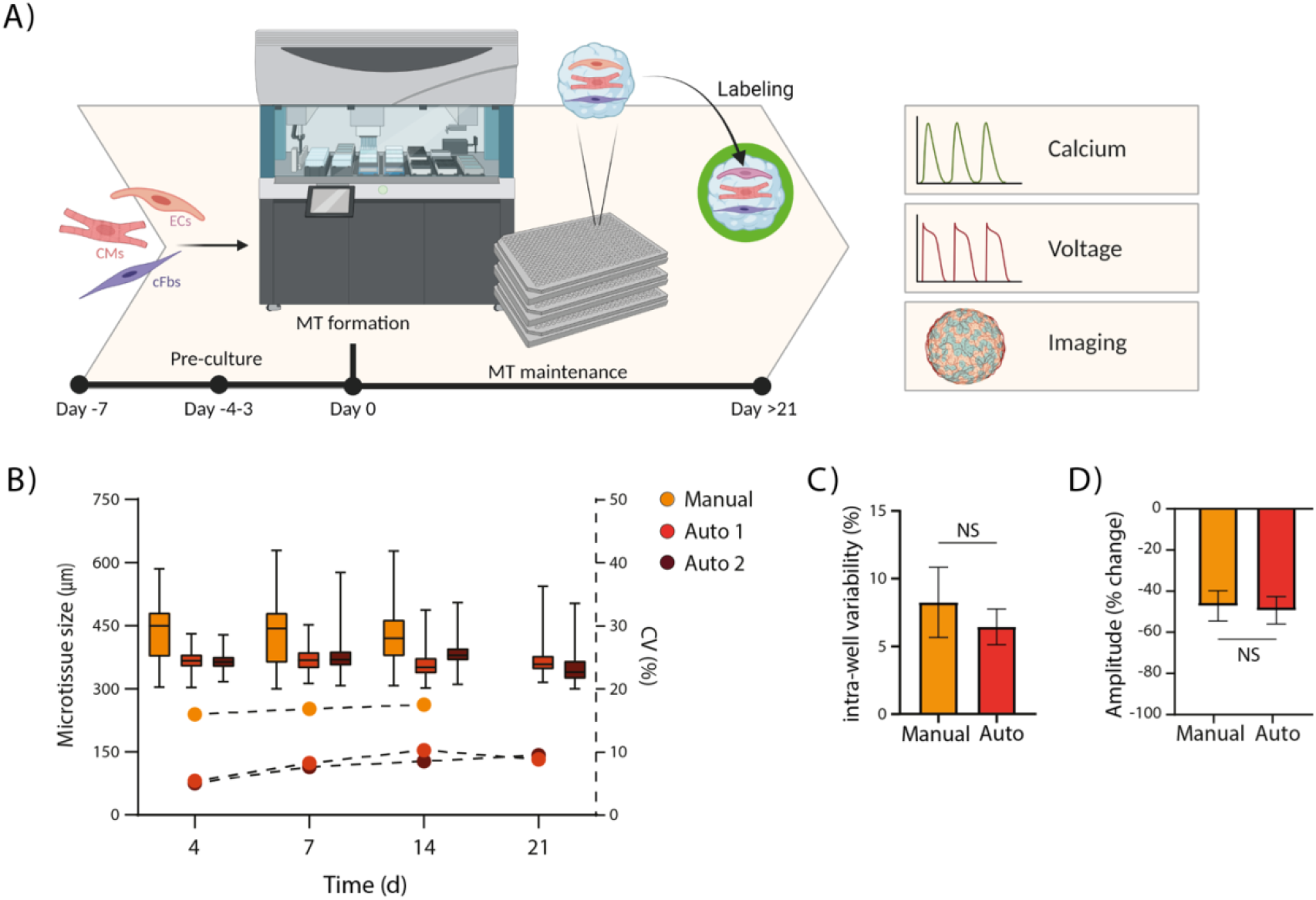
Automating cMT Generation and Functional Assays Using Robotics. **A)** Schematic overview of the automation process, including timeline and optional downstream applications. **B)** Box plots showing cMT size (left Y-axis) and a line graph displaying the corresponding coefficient of variation (CV, right Y-axis) at different days in culture. Data are shown for manually seeded cMTs (Manual, n=59-74) and cMTs generated through robotic seeding in 2 independent experiments (Auto 1, n=363-383; Auto 2, n=360-378). **C)** Intra-well variability of Ca^2+^ transients (peak-to-peak) recorded from manual-, and automated-seeded cMTs, generated on the same day and from the same cell batch. Data is mean ± SEM; n=18 (manual) and n=15 (auto); ns, not significant (unpaired t-test). **D)** Mean percentage change in Ca^2+^ peak amplitude of manual- and automated-seeded cMTs following addition of 0.75 µM nifedipine, and corrected for vehicle (DMSO). Data is mean ± SEM; n=6 for both auto and manual cMTs; ns, not significant (unpaired t-test).

A notable advantage of automation was the reduced time required for cMT generation. Seeding three 384-well plates was completed within 7 min, compared to ∼75 min needed for manual seeding and transfer. Automated dye labelling also can reduce time and variability associated with manual labelling for high-throughput imaging or screening assays. We therefore automated the labelling and washing steps for optically measuring Ca^2+^ transients and voltage kinetics. Consistent labelling across an entire 384-well plate was possible, with compound-induced autofluorescence clearly detectable (**Supplementary Fig. 2c**). Overall, transitioning to automated processes not only improved the efficiency of cMT production but also enhanced the reliability and reproducibility of functional assays.

### Cardiac microtissues show high drug response predictivity in an automated, blinded high throughput screen

We have previously demonstrated that hiPSC-CMs in 3D cMTs exhibit improved structure and metabolism compared to when in 2D monocultures^7^. Therefore, we investigated whether cMTs could also enhance the predictivity of hiPSC-CM responses to inotropic drugs, particularly in a HTS setup. Detection of positive inotropes (drugs that increase contraction force) typically require hiPSC-CMs with better myofilament organization and Ca^2+^ handling than what is obtained in 2D cultures^11,30^. Inotropic compounds primarily affect intracellular Ca^2+^ levels, either directly (e.g., through L-type Ca^2+^ channels) or indirectly (e.g., via the β-adrenergic receptor-cAMP-PKA pathway). Consequently, the majority of positive inotropic effects manifest as increases in Ca^2+^ transient peak amplitude, kinetics (rising and falling slopes), beat rate (positive chronotropy), and/or an increase in intracellular Ca^2+^ load, that is often reflected by an increase in the area under the curve (AUC) and PWD (**Supplementary Fig. 2d**).

In cMTs, the positive inotropes isoproterenol and levosimendan were observed to increase both the beat rate and Ca^2+^ transient peak amplitude, along with increases in the rising and falling slopes (**Fig. 3a**). In contrast, in 2D multi-cell type cultures generated in parallel from the same cell batches, these compounds only induced positive chronotropic responses without affecting the amplitude or kinetics. This aligns with previous findings that 2D models frequently fail to capture certain inotropic responses, particularly when the drug’s mode of action is mediated through cAMP signalling^31^.

**Figure 3.**
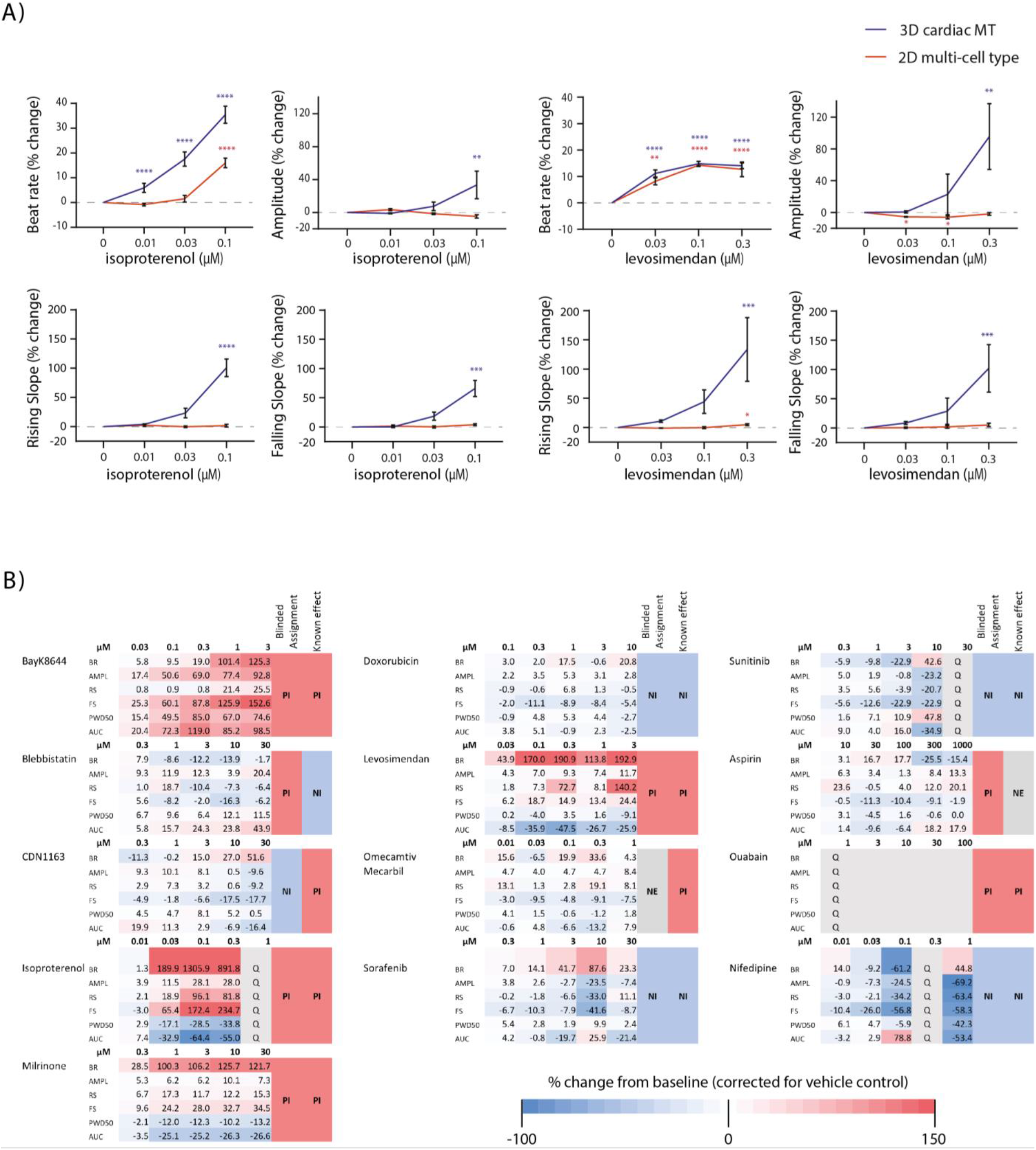
Accuracy of Drug Responses in cMTs. **A and B)** Concentration-response plots of isoproterenol (**A**) and levosimendan (**B**) on various Ca^2+^ transient parameters measured in either cMTs or 2D multi-cell type cultures generated from the same batches of cells. Data show mean percentage change from baseline, corrected for vehicle (DMSO); error bars ±SEM; n=4 (cMTs) and n=3 (2D multi-cell type) per concentration. * p<0.04, ** p<0.006, *** p<0.001, **** p<0.0001 (two-way ANOVA for global significance, followed by Sidak’s post-hoc test for multiple comparisons). **C)** Heatmap summarizing the results of a 13-reference compound blind screen performed using automated processes on cMTs, and the resulting classification of each compound as a positive inotrope (PI), negative inotrope (NI) or having no inotropic effect (NE). Data show mean percentage change from baseline, corrected for vehicle (DMSO); n=5 cMTs per concentration at baseline. BR, beat rate; AMPL, Ca^2+^ peak amplitude; RS, rising slope; FS, falling slope; PWD50, peak width duration at 50% of decay; AUC, area under the curve; Q, quiescence.

Leveraging our robotics platform, we conducted blinded screens of reference inotropic drugs, ranging from typically easily detectable compounds (e.g., L-type Ca^2+^ channel modulators), to more challenging drugs like phosphodiesterase 3 (PDE3) inhibitors. To assess reproducibility, the screen was performed using cMTs labelled with a Ca^2+^ or voltage-sensitive dye and prepared via both manual and automated seeding methods. Drug effects were classified based on overall changes in the aforementioned Ca^2+^-related parameters after exposing the cMTs to varying concentrations of these compounds (**Fig. 3b, Supplementary Fig. 3a**). Additionally, voltage kinetics and triangulation ratio (APD_30_/APD_90_) were calculated from voltage waveforms to determine whether electrophysiology was altered (**Supplementary Fig. 2d, Supplementary Fig. 3a**).

Across 3 independent screens, an accuracy rate of >80% in classifying the drugs as positive, negative, or neutral inotropes was achieved (**Supplementary Fig. 2e**). Screens utilizing cMTs from automated seeding demonstrated 100% reproducibility, with consistent compound classifications across both screens (**Supplementary Fig. 2f**). These screens also showed a 90.9% similarity with the screen conducted using manually seeded cMTs. Notably, in all screens ouabain induced atypical Ca^2+^ transients with steep rising slopes, increased peak amplitudes, and prolonged relaxation times, reducing the beat rate to levels that precluded quantification (**Supplementary Fig. 3b**). However, these visual changes in amplitude and kinetics allowed us to correctly classify ouabain as a positive inotrope.

Overall, these results underscore the capability of cMTs to accurately capture both negative and positive inotropic responses, highlighting their potential for high-throughput cardiac safety assessments.

### Efficacy screen identifies compounds that rescue arrhythmias in CPVT1 cardiac microtissues

Beyond cardiac safety assessment, predictive and automated cMT assays provide opportunities for large-scale efficacy screenings to identify compounds with therapeutic potential. We evaluated the suitability of cMTs for modelling arrhythmogenic diseases, specifically CPVT1. The cMTs were generated using hiPSC-CMs genetically engineered to harbour the CPVT1-associated RYR2 mutation p.M4019R (**Supplementary Fig. 4**). Given that β-adrenergic stimulation alone often does not reliably induce robust arrhythmogenic phenotypes *in vitro*^19,20,32^, we investigated whether bypassing β-adrenergic receptor stimulation and directly targeting the PKA pathway by treating the CPVT1-cMTs with dbcAMP would result in more consistent triggering of arrhythmias.

At baseline, regular Ca^2+^ oscillations were observed in >85% of the CPVT1-cMTs and in 100% of the isogenic control (WT-cMTs) (**Fig. 4a, b**). After 3 days of dbcAMP treatment, arrhythmias were reproducibly induced in 70.2±12.6% of the CPVT1-cMTs, compared to 17.4±15.6% of the WT-cMTs (**Fig. 4a, b**). Arrhythmic events were automatically scored using custom-developed software that facilitated high-throughput analysis and eliminated operator biases. Ca^2+^ transients were classified as either “non-arrhythmic-like”, “arrhythmic-like (Type A)”, “fibrillation-like (Type B)” or “noise” (**Supplementary Fig. 5a**).

**Figure 4.**
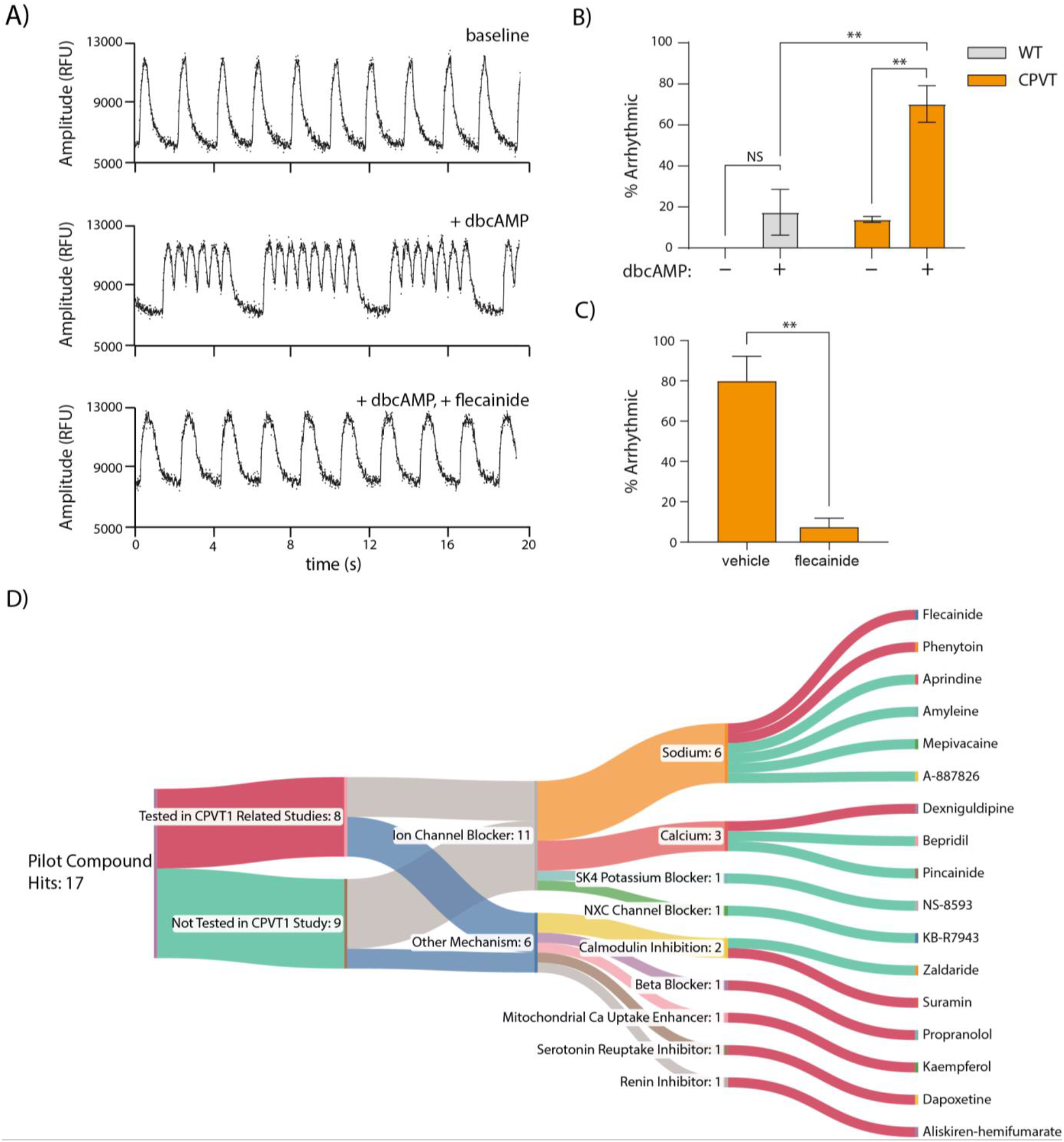
Identification of Compounds that Resolve Arrhythmias in CPVT1-cMTs. **A)** Representative Ca^2+^ transients recorded from CPVT1-cMTs under three conditions: vehicle control (baseline; non-arrhythmic), after dbcAMP treatment (+dbcAMP; arrhythmic), and after subsequent addition of flecainide (+flec; non-arrhythmic). **B)** Percentages of WT-cMTs and CPVT1-cMTs treated either with vehicle (culture medium) or 0.5 mM dbcAMP with arrhythmic phenotypes. Data show mean ±SEM; n=38 (WT +vehicle), n=53 (WT +dbcAMP), n=29 (CPVT1 +vehicle), and n=55 (CPVT1 +dbcAMP) from 2 independent experiments. ** p<0.01 (two-way ANOVA followed by Fisher’s LSD test). **C)** Percentages of dbcAMP-treated CPVT1-cMTs with arrhythmic phenotypes following subsequent addition of either vehicle (DMSO) or flecainide. Data show mean ±SEM; n=453 (vehicle) and n=457 (flecainide) from 3 independent experiments. ** p<0.006 (unpaired t-test). **D)** Sankey plot depicting the 17 hit compounds and their modes of action identified from an efficacy screen performed on arrhythmic CPVT1-cMTs. A hit is defined as a compound that achieved 100% arrhythmia rescue at 1 or more test concentrations.

In an initial screen using the automated pipeline, we assessed whether the arrhythmic events detected in the CPVT1-cMTs were related to the RYR2 mutation by treating cMTs with flecainide. Automated scoring across 3 independent experiments revealed that 92.6±6.4% of the arrhythmic CPVT1-cMTs were rescued following acute (15 min) flecainide treatment. In contrast, arrhythmias were only resolved in 20.2±17.5% of the CPVT1-cMTs treated with the vehicle control (**Fig. 4a, c**). No rescue was observed in arrhythmic WT-cMTs treated with flecainide, with instead all non-arrhythmic WT-cMTs also becoming arrhythmic (data not shown). These results reflect the protective effect of flecainide against CPVT1-induced arrhythmias, along with its known proarrhythmic risks due to its ability to inhibit IK_r_/hERG^33^.

Subsequently, in a larger screen of 92 FDA-approved drugs, dbcAMP induced arrhythmias in 100% of CPVT1 cMTs at baseline, with 17 compounds identified that completely rescued the arrhythmogenic phenotype at one or more test concentrations (**Fig. 4d**). Of these, 8 had been previously evaluated in CPVT1 studies, while 9 demonstrated novel antiarrhythmic effects in this model. Notably, 11 compounds were antiarrhythmic drugs classified as ion channel blockers, including sodium (flecainide, phenytoin, aprindine, amyleine, mepivacaine, A-887826) and calcium channel agonists (dexniguldipine, bepridil, pincainide), a SK4 potassium blocker (NS-8593), and a NXC channel blocker (KB-R7943). The remaining hits identified had unique modes of action, such as inhibiting calmodulin or serotonin transport.

## Discussion

Developing scalable HTS approaches for 3D human stem cell models remains a major challenge for disease modelling and drug discovery. In this study, we industrialized 3D cMTs for HTS to identify new and safe therapeutics by optimizing differentiation protocols for various cardiac cell types, developing fluorescence-based assays to assess Ca^2+^ and voltage in high-throughput, and confirmed the functionality and drug responsiveness of cMTs in these assays. As a final industrialization step, we employed robotics for cMT formation, maintenance and assay execution, achieving en masse production with high reproducibility.

Automated cMTs were utilized to evaluate cardiac safety in blinded screens, demonstrating high accuracy (>80% correct classification) and reproducibility (>90% across 3 independent experiments) in detecting inotropic effects. Crucially, several drugs often misclassified in 2D hiPSC-CM models, such as the Ca^2+^ sensitizer/PDE3 inhibitor levosimendan^31^, were accurately predicted in the cMT model possibly due to improved hiPSC-CM maturation or enhanced PDE3 expression due to the presence of cFbs^34^. However, the sarcoplasmic reticulum (SR) Ca^2+^-handling properties of hiPSC-CMs may still be relatively immature in cMTs, as evidenced by the inability to detect CDN1163, a SERCA activator, as a positive inotrope. Additionally, aspirin was misclassified as a positive inotrope at the highest concentrations tested (300 and 1000 µM, ≥14-fold the free C_max_) based on its effects on Ca^2+^ transient peak amplitude and rising slope. These concentrations however exceed the commonly tested range in hiPSC-cardiac models (0.1-100 µM^5,35,36^), where we did not observe such effects. Furthermore, unlike with EHTs^6,37,38^, we are unable to directly measure contraction force in cMTs, leading to the misclassification of drugs like blebbistatin, a myosin II inhibitor.

We also established a robust arrhythmogenic assay for inherited cardiac arrhythmic disorders, such as CPVT1, that is amenable for high-throughput. This is significant as most previous CPVT1 hiPSC-CM models have relied on low-throughput 2D models and outputs with relatively low reproducibility^19,20,39,40^. For example, Fatima *et al*. and Kujala *et al*. reported arrhythmic activity rates of 32% and 30% respectively in CPVT1-CMs after adrenergic stimulation using electrophysiological assays^19,20^. Additionally, Novak *et al*. noted that only 16% of CPVT1 hiPSC-CMs showed arrhythmic behaviour in Ca^2+^ transients after isoproterenol treatment^32^.

In contrast, we could reproducibly trigger arrhythmias in >70% of CPVT1-cMTs by treating them with dbcAMP for 3 days. We hypothesize that this occurs because dbcAMP bypasses adrenergic stimulation, raising intracellular cAMP to levels much higher than typically achieved with β-adrenergic stimulation alone. This may be due to the generally low basal receptor tone and low endogenous cAMP in hiPSC-CMs^41^. These elevated intracellular cAMP levels subsequently increase PKA activity and excessive Ca^2+^ release from the SR through dysfunctional RYR2 channels, which in turn causes Ca^2+^ overload and initiates arrhythmias.

To validate our efficacy screen, we treated dbcAMP-induced arrhythmic CPVT1-cMTs with flecainide, a known antiarrhythmic agent^24^. We demonstrated that acute flecainide treatment effectively rescued the arrhythmic events, likely through a dual mode of action involving the suppression of spontaneous SR Ca^2+^ release events via RYR2 inhibition and suppression of triggered beats via Nav1.5 channel block^24^. The standard treatment for CPVT1 patients involves the use of β-blockers, with flecainide included in cases where additional suppression of arrhythmic episodes is required^42^. Despite this, ∼10-25% of patients do not respond adequately^25^. Identifying novel therapies therefore is crucial to provide effective treatment options for non-responders and to improve the overall management of CPVT1.

As a proof-of-concept, we screened 92 FDA-approved drugs. These included compounds with potential antiarrhythmic effects (e.g. ion channel blockers), no expected antiarrhythmic effects (e.g. aspirin), agents that might exacerbate the arrhythmias (e.g. caffeine), and drugs with unclear but potentially relevant antiarrhythmic modes of action (e.g. suramin). Of the 17 compounds that completely rescued the arrhythmogenic phenotype at one or more test concentrations, the majority (65%) were ion channel blockers, most likely acting by reducing Ca^2+^ overload in the arrhythmogenic hiPSC-CMs. Interestingly, dapoxetine, a selective serotonin reuptake inhibitor (SSRI), was identified as a hit, consistent with recent findings suggesting SSRIs may be a therapeutic option for CPVT1 patients^43,44^. Notably, 9 of the 17 compound hits had not been previously tested for treating CPVT1, highlighting the potential of this HTS assay to identify novel therapeutic agents. Among these was zaldaride, an antidiarrheal agent that inhibits calmodulin^45^. Given that mutations in calmodulin genes are known to cause cardiac arrhythmic diseases and that calmodulin interacts with RYR2 and the L-type calcium channel^46,47^, the antiarrhythmic mechanism of zaldaride may involve inhibiting these interactions.

Interestingly, dantrolene, a drug effective for malignant hyperthermia and previously shown to rescue the arrhythmic phenotype in CPVT1-CMs generated from patient hiPSC lines^18,48^, did not show a significant antiarrhythmic effect in our screen. This is possibly due to the genetic background of the hiPSC line or the *RYR2* mutation. The RYR2 mutation p.S406L is thought to act via a “domain unzipping” mechanism, with dantrolene believed to stabilise the interdomain interactions^49,50^. In contrast, the p.M4019R mutation studied here is in the cytoplasmic domain but much closer to the SR transmembrane domain, potentially altering its responsiveness to dantrolene. Follow-up assays will be required to confirm hits and compare their effects in CPVT1 cMTs containing hiPSC-CMs with different *RYR2* mutations or derived from CPVT patients with different arrhythmogenic phenotypes to evaluate varying drug responsiveness. If these features can be reliably captured and distinguished in cMTs, it could provide new opportunities to stratify patients and personalize their treatments.

In conclusion, automating the formation and maintenance of cMTs, as well as HTS assays for both safety and efficacy assessments, offers several advantages. Namely, reduced variability and enhanced cardiomyocyte maturation within the cMTs, and reliable drug screening with high reproducibility. This approach holds significant promise for further HTS applications and identifying potential novel therapeutic targets for inherited cardiac arrhythmic diseases like CPVT1.

