## Supplementary Figures for "Industrialization of three-dimensional hiPSC-cardiac microtissues for high-throughput cardiac safety and drug discovery screening"

**Supplementary Figure 1. Characterization of hiPSC-derived cardiac endothelial cells, cardiomyocytes and cardiac fibroblasts**

**Supplementary Figure 2. Evaluating automated processes for cMT formation and functional screening**

**Supplementary Figure 3. Effects of compounds on $Ca^{2+}$ and voltage parameters in cMTs**

**Supplementary Figure 4. Generation of hiPSC line with *RYR2* mutation**

**Supplementary Figure 5. Automated analysis of $Ca^{2+}$ traces for arrhythmias**

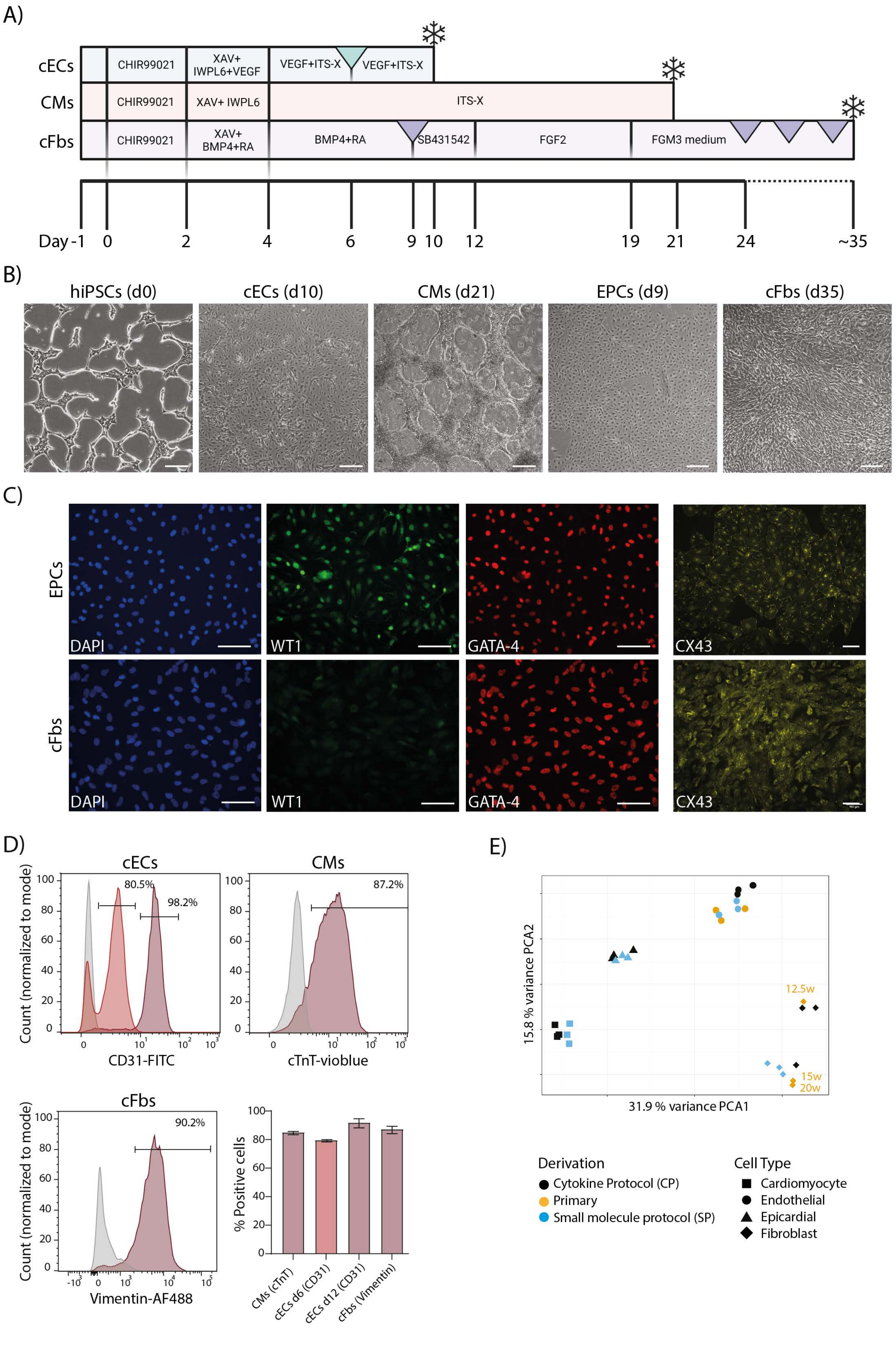
A)
cECs
CMs
cFbs
CHIR99021
XAV+ IWPL6+VEGF
VEGF+ITS-X
VEGF+ITS-X
XAV+ IWPL6
ITS-X
XAV+ BMP4+RA
BMP4+RA
SB431542
FGF2
FGM3 medium
Day -1
0
2
4
6
9
10
12
19
21
24
~35
B)
hiPSCs (d0)
cECs (d10)
CMs (d21)
EPCs (d9)
cFbs (d35)
C)
EPCs
cFbs
DAPI
WT1
GATA-4
CX43
D)
cECs
80.5%
98.2%
Count (normalized to mode)
CD31-FITC
CMs
87.2%
cTnT-vioblue
cFbs
90.2%
Vimentin-AF488
% Positive cells
CMs (cTnT)
cECs d6 (CD31)
cECs d12 (CD31)
cFbs (Vimentin)
E)
15.8 % variance PCA2
31.9 % variance PCA1
12.5w
15w
20w
Derivation
Cytokine Protocol (CP)
Primary
Small molecule protocol (SP)
Cell Type
Cardiomyocyte
Endothelial
Epicardial
Fibroblast

***Supplementary Figure 1. Characterization of hiPSC-derived cardiac endothelial cells, cardiomyocytes and cardiac fibroblasts.***

***A)*** *Schematic of protocols for differentiating hiPSCs into cardiac endothelial cells (cECs), cardiomyocytes (CMs), and cardiac fibroblasts (cFbs). The timeline indicates the timepoints for the addition and removal of cytokines and small molecules, as well as passaging (triangle) and freezing (snowflake) the differentiated cells.*

***B)*** *Representative phase contrast images of hiPSCs at day 0 and the differentiated cell types at the end of their respective differentiation protocols (d, days of differentiation). Scale bar: 100 µm.*

***C)*** *Immunofluorescence images showing the expression of WT1, GATA4, and CX43 in epicardial cells (EPCs), and the subsequent cFbs. Scale bar: 100 µm.*

***D)*** *Flow cytometry analysis with representative histograms showing the expression of the EC marker CD31 directly after isolation (day 6) and before cryopreservation (day 12), the CM marker cardiac troponin-T (cTnT), and the cFb marker vimentin. Isotype controls are shown in gray, with values indicating the percentage of cells positive for each marker. The bar graph (bottom right) shows the mean percentages ±SD of cells positive for each marker in the respective cell types (n=3 differentiations).*

***E)*** *Principal-component analysis (PCA) of RNA-seq profiles from CMs, cECs, EPCs and cFbs differentiated from the hiPSC line LUMC0020iCTRL-06 using the "small molecule" protocols described in (A), compared to samples obtained using cytokine-based protocols (CP) and primary human material. Analysis was performed using genes filtered based on their expression across all samples (CMs, 18336 genes; cECs, 17126 genes; EPCs, 16857 genes; cFbs, 19684 genes). Symbols represent individual samples.*

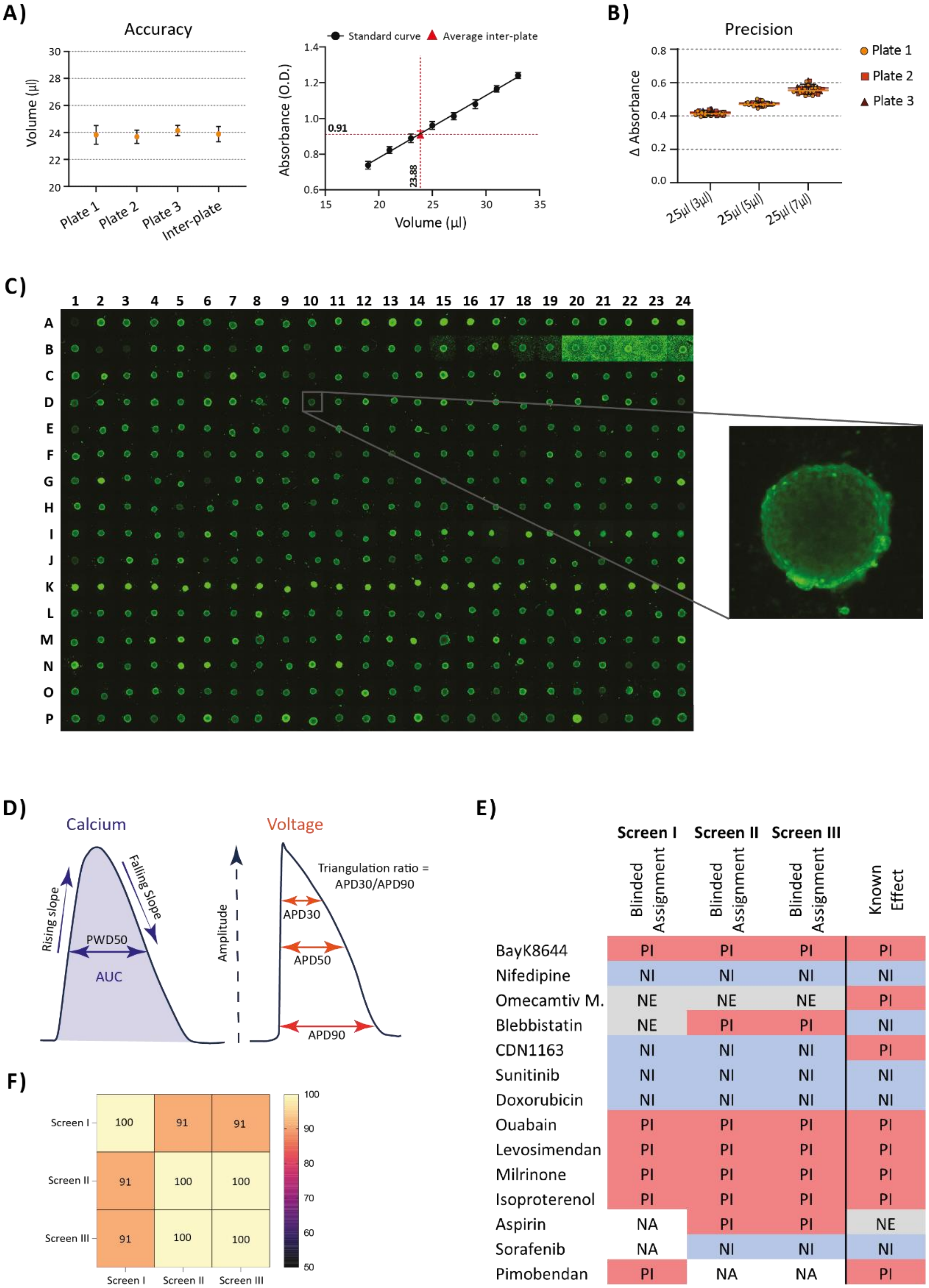

A)
Accuracy
Volume (µl)
30
28
26
24
22
20
Plate 1
Plate 2
Plate 3
Inter-plate
Standard curve
Average inter-plate
Absorbance (O.D.)
1.4
1.2
1.0
0.8
0.6
0.91
23.88
15
20
25
30
35
Volume (µl)
B)
Precision
Δ Absorbance
0.8
0.6
0.4
0.2
0.0
25µl (3µl)
25µl (5µl)
25µl (7µl)
Plate 1
Plate 2
Plate 3
C)
1 2 3 4 5 6 7 8 9 10 11 12 13 14 15 16 17 18 19 20 21 22 23 24
A B C D E F G H I J K L M N O P
D)
Calcium
Rising slope
Falling Slope
PWD50
AUC
Voltage
Amplitude
Triangulation ratio = APD30/APD90
APD30
APD50
APD90
E)
Screen I
Screen II
Screen III
Blinded Assignment
Blinded Assignment
Blinded Assignment
Known Effect
BayK8644 PI PI PI PI
Nifedipine NI NI NI NI
Omecamtiv M. NE NE NE PI
Blebbistatin NE PI PI NI
CDN1163 NI NI NI PI
Sunitinib NI NI NI NI
Doxorubicin NI NI NI NI
Ouabain PI PI PI PI
Levosimendan PI PI PI PI
Milrinone PI PI PI PI
Isoproterenol PI PI PI PI
Aspirin NA PI PI NE
Sorafenib NA NI NI NI
Pimobendan PI NA NA PI
F)
Screen I 100 91 91
Screen II 91 100 100
Screen III 91 100 100
Screen I
Screen II
Screen III
100
90
80
70
60
50

***Supplementary Figure 2. Evaluating automated processes for cMT formation and functional screening***

***A)*** *Evaluation of pipetting volume accuracy of the liquid handling robot. Graph (left) shows mean ± SD of the pipetted volume for each plate individually (n=384 wells) and for all plates combined (N=3 plates). Volume was determined by measuring the absorbance of the tartrazine dye and comparing to the standard curve (right).*

***B)*** *Evaluation of the precision of the liquid handling robot in pipetting small volumes (value in brackets) within a total volume of 25 µl. Graph shows the mean change (D) in absorbance for each plate, including individual datapoints (N=3 plates).*

***C)*** *Images captured of cMTs formed using a liquid-handling robot in a 384-well plate and loaded with Calcium-6 dye (green). Images were taken post-compound addition.*

***D & E)*** *Reference $Ca^{2+}$ and voltage transients (**D**) illustrating the relevant parameters (excluding beat rate) used as surrogate readouts for classifying compounds as positive inotropes (PI), negative inotropes (NI), or non-inotropic effect (NE) compounds, and summary table (**E**) of three independent blinded screens (12-13 reference compounds per screen), with 6 different $Ca^{2+}$-related parameters used for classification. The myosin modulators, omecamtiv mecarbil ("Omecamtiv M.") and blebbistatin, are expected to be classified as "NE" in a $Ca^{2+}$ assay. NA, not applicable.*

***F)*** *Heat plot reflecting the overlap in accuracy (as a percentage) between the three independent screens.*

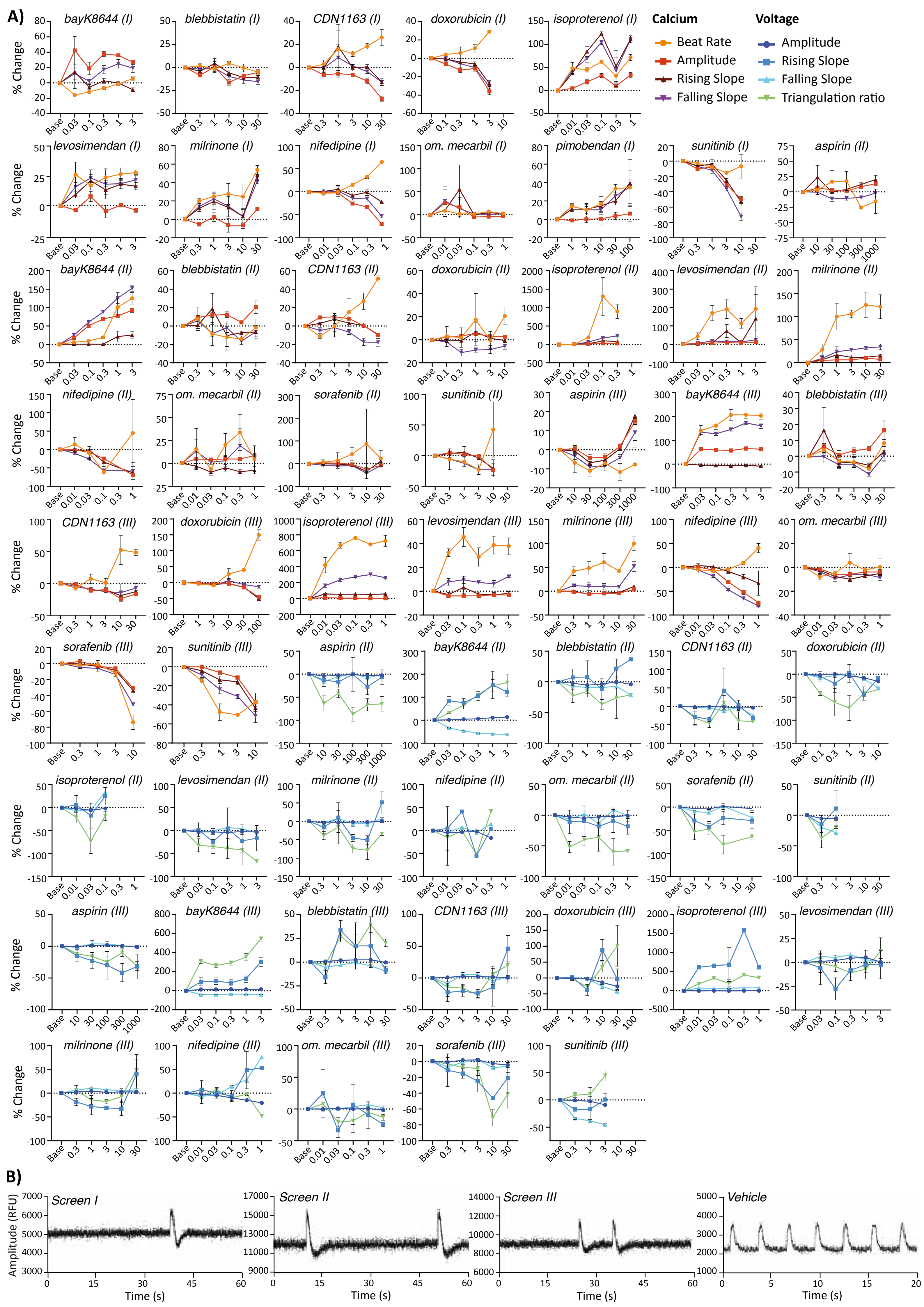
A)
Calcium
Beat Rate
Amplitude
Rising Slope
Falling Slope
Voltage
Amplitude
Rising Slope
Falling Slope
Triangulation ratio
% Change
bayK8644 (I)
blebbistatin (I)
CDN1163 (I)
doxorubicin (I)
isoproterenol (I)
levosimendan (I)
milrinone (I)
nifedipine (I)
om. mecarbil (I)
pimobendan (I)
sunitinib (I)
aspirin (II)
bayK8644 (II)
blebbistatin (II)
CDN1163 (II)
doxorubicin (II)
isoproterenol (II)
levosimendan (II)
milrinone (II)
nifedipine (II)
om. mecarbil (II)
sorafenib (II)
sunitinib (II)
aspirin (III)
bayK8644 (III)
blebbistatin (III)
CDN1163 (III)
doxorubicin (III)
isoproterenol (III)
levosimendan (III)
milrinone (III)
nifedipine (III)
om. mecarbil (III)
sorafenib (III)
sunitinib (III)
aspirin (II)
bayK8644 (II)
blebbistatin (II)
CDN1163 (II)
doxorubicin (II)
isoproterenol (II)
levosimendan (II)
milrinone (II)
nifedipine (II)
om. mecarbil (II)
sorafenib (II)
sunitinib (II)
aspirin (III)
bayK8644 (III)
blebbistatin (III)
CDN1163 (III)
doxorubicin (III)
isoproterenol (III)
levosimendan (III)
milrinone (III)
nifedipine (III)
om. mecarbil (III)
sorafenib (III)
sunitinib (III)
Base
B)
Screen I
Screen II
Screen III
Vehicle
Amplitude (RFU)
Time (s)

***Supplementary Figure 3. Effects of compounds on $Ca^{2+}$ and voltage parameters in cMTs***

***A)*** *Line graphs showing the results from three independent blinded $Ca^{2+}$ and voltage screens (I, II and III), with 12-13 compounds evaluated per screen, 5 concentrations per compound, n=5 cMTs per concentration at baseline. For each compound concentration tested, the percentage change compared to baseline (corrected for vehicle) is plotted for various $Ca^{2+}$ and voltage parameters ($Ca^{2+}$: beat rate, peak amplitude, rising slope, falling slope; voltage: peak amplitude, rising slope, falling slope, triangulation ratio).*

***B)*** *Representative $Ca^{2+}$ traces from the three independent blinded screens (I, II and III) illustrating the effect of ouabain on cMTs, alongside a vehicle control (0.1% DMSO) trace (bottom right). RFU, relative fluorescence unit.*

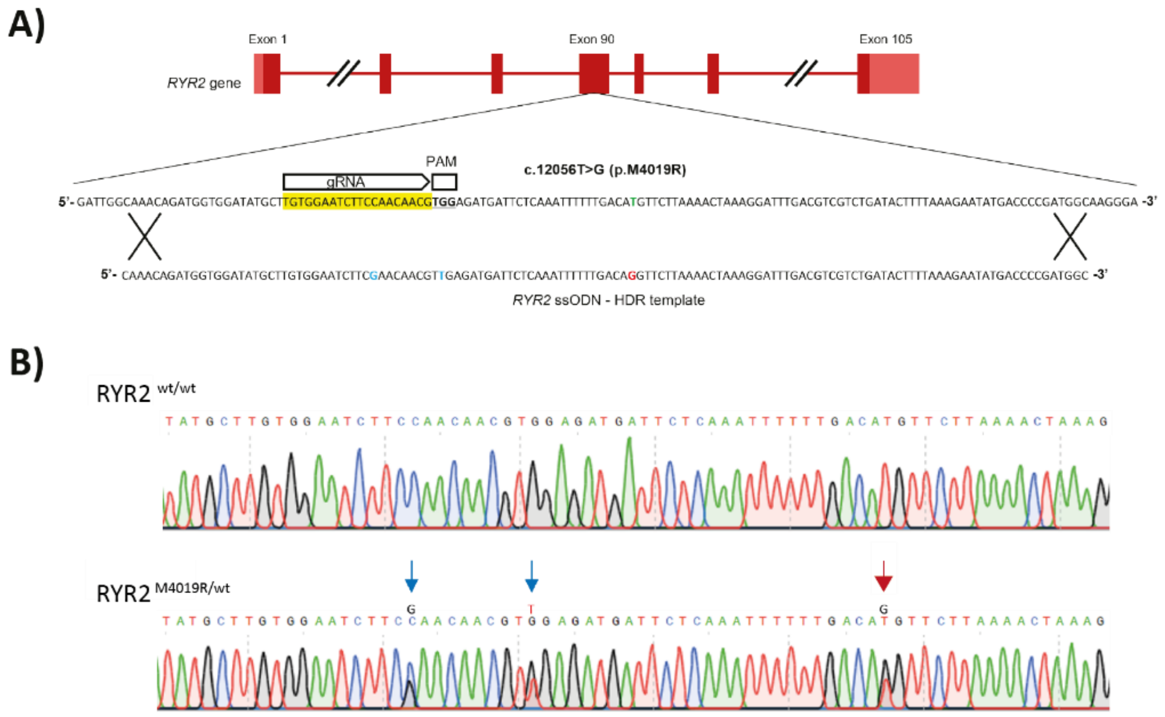


***Supplementary Figure 4. Generation of hiPSC line with RYR2 mutation***

***A)*** *Schematic outlining the strategy to introduce the RYR2 c.12056T>G (p.M4019R) mutation by homologous recombination into exon 90 of the RYR2 wild-type sequence. The guide RNA (gRNA) and corresponding protospacer adjacent motif (PAM) sequences are indicated, along with the sequence of the single-stranded oligodeoxynucleotide (ssODN) used to introduce mutation (red). Silent mutations used to assist with targeting and screening are indicated in blue.*

***B)*** *Sequence analysis of PCR-amplified RYR2 exon 90 from the CTRL ($RYR2^{wt/wt}$) and the CPVT1 ($RYR2^{M4019R/wt}$) hiPSC lines. The red arrow indicates the heterozygous CPVT1-associated mutation and the blue arrows indicate the introduced silent mutations.*

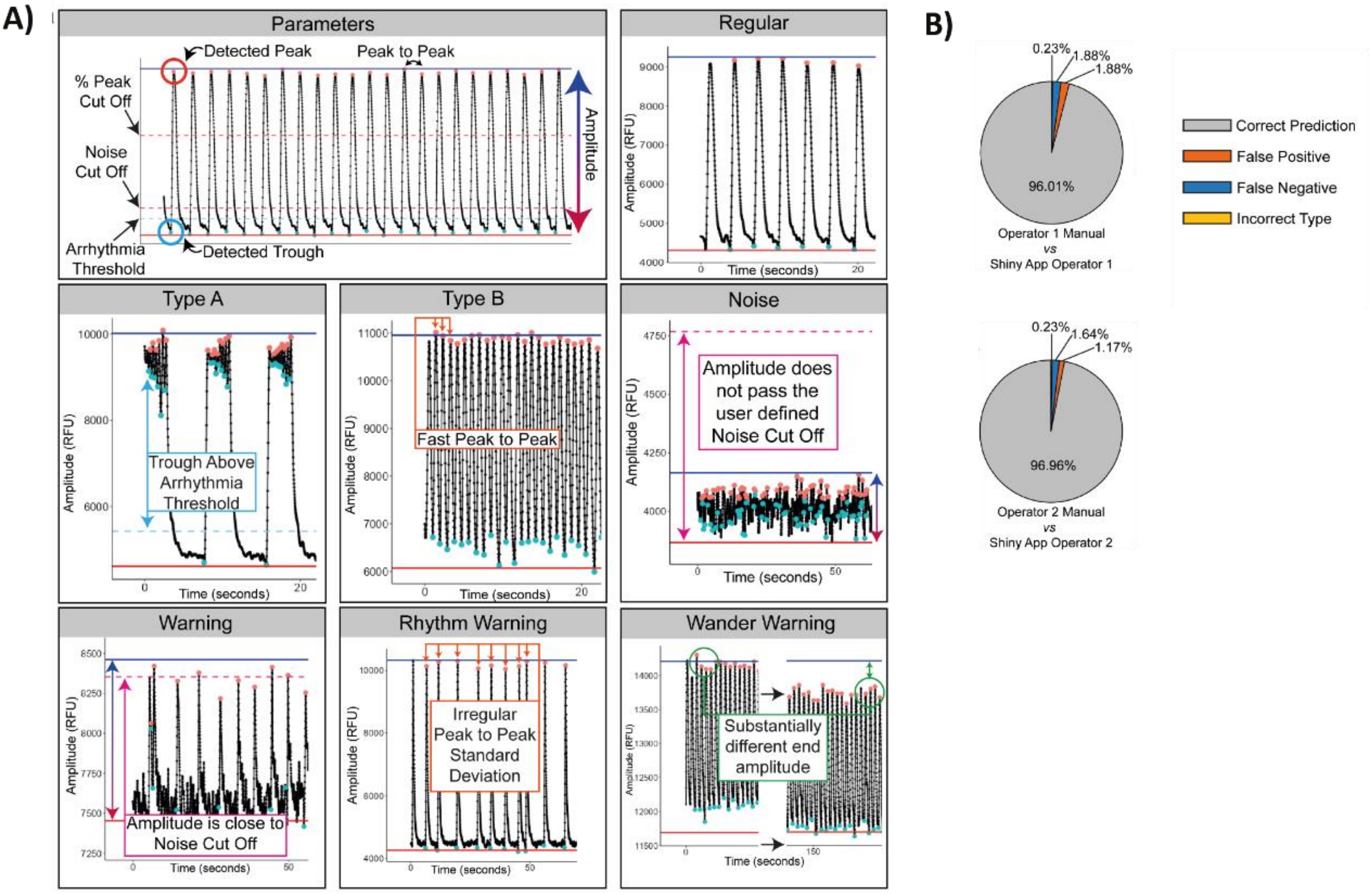


***Supplementary Figure 5. Automated analysis of $Ca^{2+}$ traces for arrhythmias***

***A)** Overview of parameters calculated based on user-defined criteria from an inputted trace (top left), and example traces for the different classifications determined by the software program. The primary reason for each non-rhythmic classification is indicated within each box.*

***B)** Pie charts indicating the percentages of traces that were classified the same or different by the software and 2 independent experts (Operator). "False positive" and "false negative" refers to percentages of traces incorrectly classified by the software as either arrhythmic or regular, respectively, compared to the classification made by the expert. "Incorrect type" indicates that both the software and expert classified the signal as arrhythmic, but the type of arrhythmia that was assigned to the trace was not matching. N=427 traces.*
